# A Novel ex-vivo platform for personalized treatment in metastatic ovarian cancer

**DOI:** 10.1101/2024.03.14.585117

**Authors:** Alain Valdivia, Adebimpe Adefolaju, Morrent Thang, Luz Andrea Cuaboy, Catherine John, Breanna Mann, Andrew Satterlee, Victoria L Bae-Jump, Shawn Hingtgen

## Abstract

The lack of functional precision models that recapitulate the pathology and structure/function relationship of advanced ovarian cancer (OC) within an appropriate anatomic setting constitutes a hurdle on the path to developing more reliable therapies and matching those therapies with the right patients. Here, we developed and characterized an Organotypic Mesentery Membrane Culture (OMMC) model as a novel ex-vivo platform where freshly resected human patient OC tumor tissue or established cell lines are seeded directly atop living intact rat mesenteric membranes, rapidly engraft, and enable functional assessment of treatment response to FDA-approved standard care of treatment as single and combination drug therapies within just five days. This study showed successful survival of dissected mesentery tissue, survival and engraftment of tumor cells and patient tumor tissue seeded on OMMCs, mesentery-tumor cell interaction, and quantification of tumor response to treatment and off-target toxicity. Summarized “drug sensitivity scores”, using a multi-parametric algorithm, were also calculated for each patient’s treatment response, enabling us to suggest the most effective therapeutic option. Finally, we compared drug sensitivity results from patient tumor tissue on OMMCs to matched outcomes of individual patients in the clinic and identified positive correlations in drug sensitivity, beginning to validate the functionality of OMMCs as a functional predictor of treatment response.

**Summary sentence:** We have successfully developed and characterized a novel ex-vivo platform for personalized treatment of metastatic ovarian cancer.

## 1. Introduction

Ovarian cancer (OC) is one of the most common gynecological cancers around the world with a 5-year survival rate of 50% of the cases diagnoses 2013-2019 (*1,2*) in US alone. Recent improvements in sensitivity and specificity of FDA-approved OC blood biomarker assays such as carcinoembryonic antigen (CEA), cancer antigen 125 (CA125), and human epididymis protein 4 (HE4) have provided a better OC stratification and important guidance for treatment, but this information is still insufficient to guide true precision oncology, as clinical and intratumoral heterogeneity, tumor recurrence, tumor drug resistance, and different levels of chemosensitivity contribute to variable therapeutic outcomes and make it difficult to effectively define the most efficacious therapy for each patient.

There is an urgent need to include functional testing of live tumor tissue in the current clinical diagnostic landscape. Direct measurements of tumor response to drugs *ex vivo* are an ongoing alternative under study. In recent years, 3D models, Spheroids (*3,4*), patient-derived xenograft (PDX) models (*5–8*), Organoids (*9–14*), tumor explants (*15,16*), and organotypic models (*17–21*) of OC have increasingly attracted the attention of scientists. In the case of tumor explants and organotypic models, freshly resected tumors, if maintained alive in the research laboratory at least for a short period, can maintain tissue architecture, spatial organization, tumor microenvironment, and even inherent intra-tumoral heterogeneity (*22,23*). This allows the evaluation of tumor cell behavior within their extracellular matrix and surrounding microenvironment which may suggest fidelity in the drug tumor response (*24*).

*In vitro* and *in vivo* models have become the gold standard for preclinical research and often do not offer enough relevant, reliable, or rapid information to directly guide clinical care. These deficiencies are most manifested in an inability to maintain resected patient tissue in a manner that allows functional analysis *ex vivo,* such as quantifying tumor cell kill after treatment. An ideal ovarian cancer model would bridge preclinical research and clinical care by (1) enabling engraftment of resected patient tumor tissue within its native anatomic environment *ex vivo*, (2) quantifying drug response of such tissue against experimental and approved therapies, (3) analyzing relative response rates of tumor and drug panels to provide insight on the most effective treatment for each patient, and (4) providing summarized readouts within a rapid timeframe to help guide treatment decisions. Such a model could improve the entire drug development pipeline.

In this study, we introduce the organotypic mesentery membrane culture (OMMC) model. This *ex vivo* platform acts as a living tissue substrate on which live, uncultured, ovarian tumor tissue resected from patients can be rapidly engrafted, treated, and analyzed to predict therapeutic outcomes in the clinic. We first characterized, optimized, and tested the OMMC platform and assay by seeding established human ovarian cancer cell lines and assessing tumor growth, mesentery-tumor interactions, tumor response to treatment with conventional FDA-approved standard-of-care treatments for OC, and off-target toxicity to the OMMC substrate itself. Later, using alive, surgically dissected OC tissue from patients at UNC Hospitals, we show tumor engraftment, survival, response to therapy on OMMCs. We also use a previously validated multipoint algorithm (*25*) to calculate a normalized, summarized, “Drug Sensitivity Score” and compare the drug-induced killing of established tumor cell lines and patient tumor tissues. Finally, we compare drug sensitivity results of resected patient tumor tissues on OMMCs to the actual clinical response of matched patients.

## Results

### OMMCs generation, characterization, and optimization

In the OMMC system, living mesenteric tissue is used as a living tissue substrate to capture fresh OC tissue and established OC cell lines. To first establish living rat mesenteric membranes, mesenteric tissue from 8-week-old rats was aseptically removed, carefully transferred into transwell membranes, and seeded in plates with media that created an air-tissue interface (Fig. 1A). The region of interest in the mesentery cultures, where tumor cells are eventually seeded, is shown in Figure 1B. The thin membrane of this desired region is circumscribed by fatty tissue which is not involved in any analysis. This thin membrane contains a net of collagen, elastin fibers, and cells, as observed by brightfield microscopy and hematoxylin & eosin staining (Fig. 1B, right). We confirmed the viability of mesenteric tissue using two different approaches. First, we transduced the mesenteries with lentiviral vectors encoding mCherry and luciferase. Luciferase-based viability readouts of the thin inner membranes taken at 4-, 6-, 8-, and 10-days post-seeding revealed robust luciferase signal with no statistical differences between days (Fig. 1C). To confirm these imaging results, we used our established nuclear permeability assay using propidium iodide (PI) as a fluorescent marker of cell death (*25*). High-resolution imaging of the PI-treated mesentery again showed robust and stable viability of the mesentery through 10 days. Interestingly, the mesenteric viability decreased slightly at later time points, reducing by ∼20% at 17 days post-seeding in our OMMC system (Fig. 1D). To validate the PI toxicity assay on OMMCs, quantitative imaging and analysis of OMMCs treated with increasing concentrations of DMSO (0-100%) detected subtle differences in viability and showed a dose-dependent increase in OMMC toxicity (Fig. 1E).

**Figure 1.**
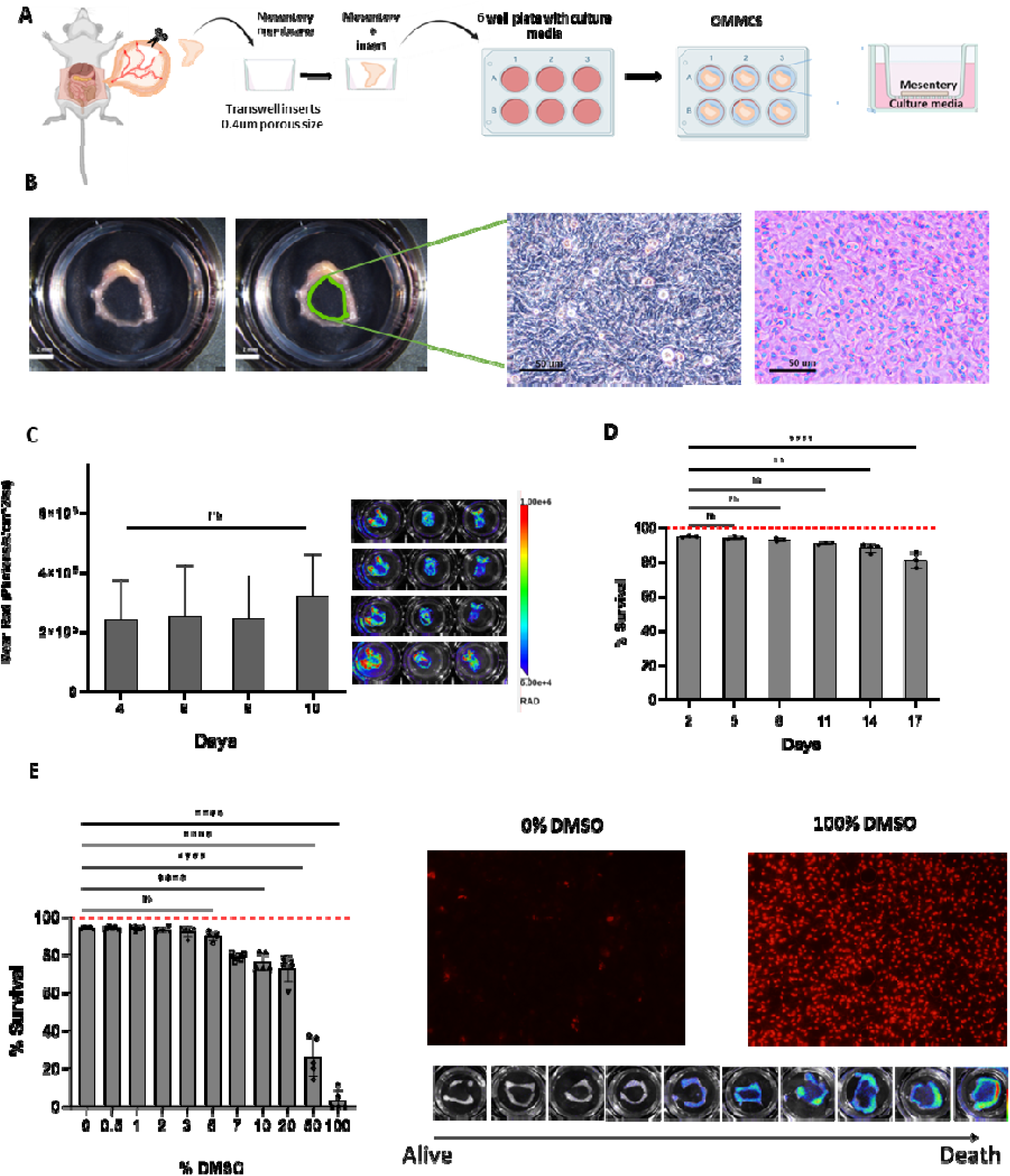
OMMC generation and mesentery viability. (A) Scheme of OMMCs generation. (B) Region of interest (green drawing) of the isolated mesentery and display of its net of cells and extracellular components by light microscopy and H&E staining. (C) BLI tracking of the transduced of 8-week-old rat mesentery over 10 days. One­way ANOVA p=0.149. (D) Confirmatory 8-week-old rat mesentery survival using the PI assay. One-way ANOVA p>0.05, **p<0.001, and ****p<0.0001. (E) Mesentery killing by increasing DMSO concentrations (E, left), one­way ANOVA p<O.OOOl. Photos of mesentery cell fluorescence taken with the AMI optical system (E, right-bottom) and inverted fluorescent scope (E, right-top), showing a gradually increasing PI signal on dead cells when exposed to 0% and 100% DMSO respectively. One-way ANOVA ****p<0.0001.

We next assessed the impact of various procedural changes on OMMC viability and cell density. First, a comparative analysis of 50 OMMCs across multiple batches showed the ability of our approach to consistently generate living mesenteric tissue (Fig. 2A). We next investigated the impact of rat age on mesenteric viability. Living mesenteric membranes were generated from rats of varying ages (3-16 weeks old) and the adult dam. As shown in Figure 2B, consistently more than 80% of the cells on these mesenteries from animals at least 8 weeks old stay viable through day 11 after dissection, followed by reduction of viability at day 14. Mesentery tissues from 3- and 4-week-old rat pups did not maintain adequate quality over time: after initially displaying high viability, spontaneous tissue compaction around day 5 caused the membranes to collapse and prevented the region of interest from being exposed or readable (Fig. 2C). We also observed an increase in fat content on mesenteric membranes which correlated with increasing rat age and resulted in smaller regions of interest (Fig. 2D); in some cases, bleeding was recorded during the isolation process. Lastly, we investigated the impact of the anatomic region on the mesenteric tissue by isolating mesenteric tissues from two main regions of the small digestive system: the ileum and jejunum (Fig. 2E). Using confocal microscopy and PI staining, quantitative imaging showed no significant difference in the cell content between the regions (∼ 600 cells/mm^2^); however, there was a small but significant difference in mesentery thickness between regions with the ileum region showing increased thickness compared to the jejunum (∼ 28 µm ileum and ∼ 22 µm Jejunum) (Fig. 2F). From these findings, we restricted our mesenteric isolation to Ileum regions dissected from 8-week-old rats.

**Figure 2.**
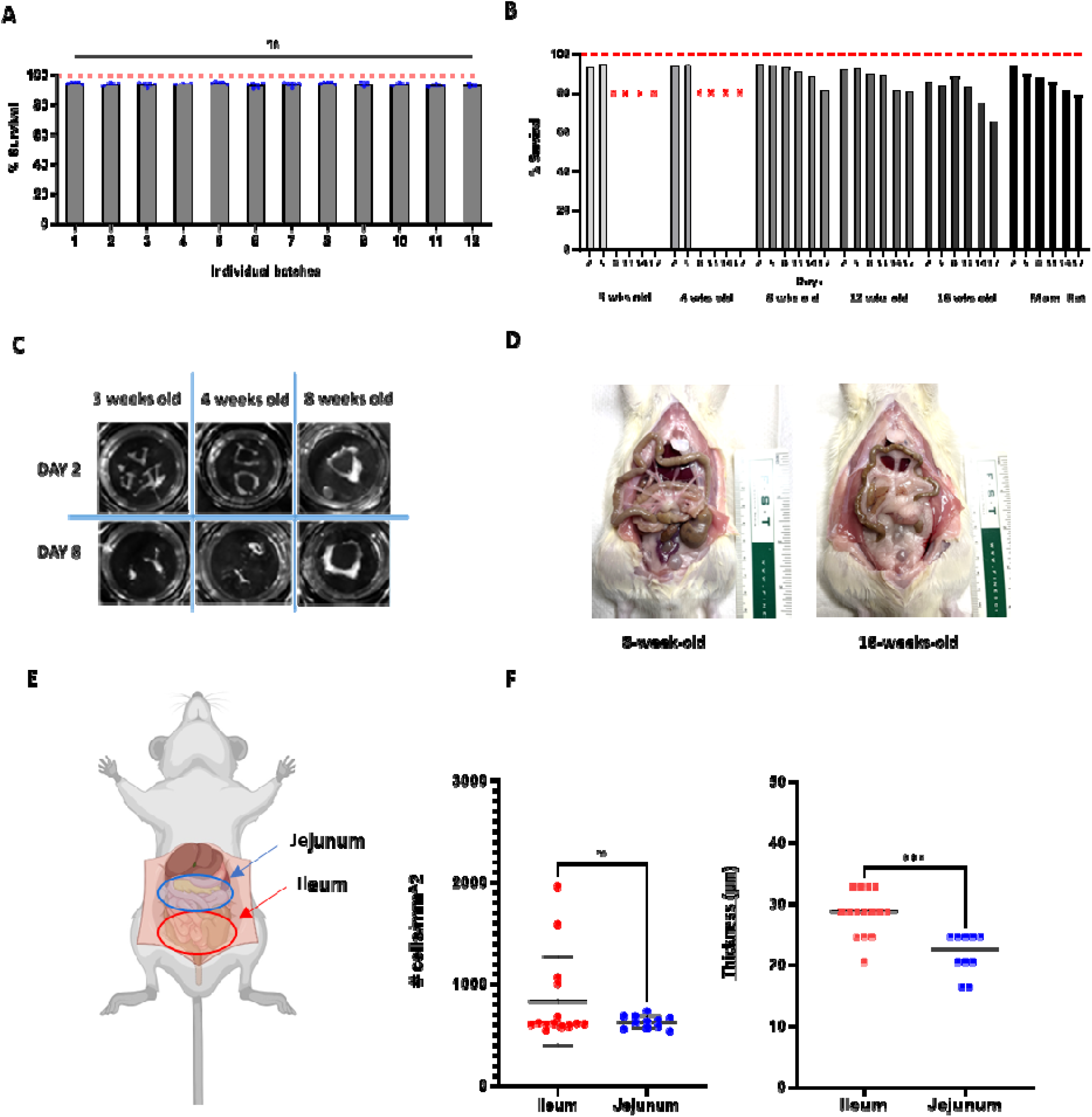
Impact of different factors during the OMMC generation process and optimization. (A) Consistent mesentery survival of 8-week-old rats observed and confirmed by PI assay across experiments over 6 days. One-way ANOVA p=0.419. (B) Mesentery survival gradually decreased from Day 11 onwards for all ages evaluated, except for 3 and 4 weeks old where the PI fluorescence was not measurable after Day 5 on (B, red crosses). (C) A photograph showed shrinkage of the region of interest in 3-and 4-week-old rat mesenteries by Day 8. (D) Real photos of the exposed mesenteries of 8-week-old and 16-week-old rats. The older rat showed higher fat content around the region of interest. (E) Cell count and membrane thickness of Ileum and Jejunum using PI staining and confocal imaging. There was a homogenous number of cells in both regions not showing a significant difference, T-test, p=0.133. However, a difference was noticed in thickness between the regions. T-test ***p<0.001.

### Establishing tumors in the OMMC system

The ultimate goal of the OMMC platform is to engraft, treat, and analyze established cell lines and human patient tumor tissues. Having established the mesenteric component of the platform, we next focused on the OC tumor component. To simplify development, we began our testing with two of the most well-established human OC cell lines: ES-2 and SKOV3. These cell lines were transduced with reporter genes mCherry-Firefly Luciferase. At Day 0, 5 to 7 tumor spots each containing ∼2×10^4^ cells were placed onto the established mesenteric tissue (Fig. 3A). Brightfield imaging was used to visualize tumor spots, and then serial fluorescence and bioluminescence imaging was used to monitor growth, expansion, and viability of each foci (Fig. 3B). Each OC foci formed a rounded and uniform tumor spot on the rat mesentery, recapitulating the metastatic tumor morphology which develops on the human omentum and mesentery in OC patients at advanced stages (*26*). Longitudinal cell viability tracking showed that both cell lines exhibited consistent growth and expansion on the mesentery, expanding 4-fold over 10 days (Fig. 3C). To confirm consistency on the seeding process, we calculated an inter-well variability of the initial OC seeding (Day 0) of 216 tumor foci and distributed on 36 individual mesenteric tissues. Results showed only 2.7% of variability on tumor seeding in between wells was observed (Fig. 3D). Comparison of growth rates showed similar consistency, as no significant differences in the viability of ES-2 or SKOV3 foci was detected 5 days post-seeding when compared across 4 individual batches (Fig. 3E).

**Figure 3.**
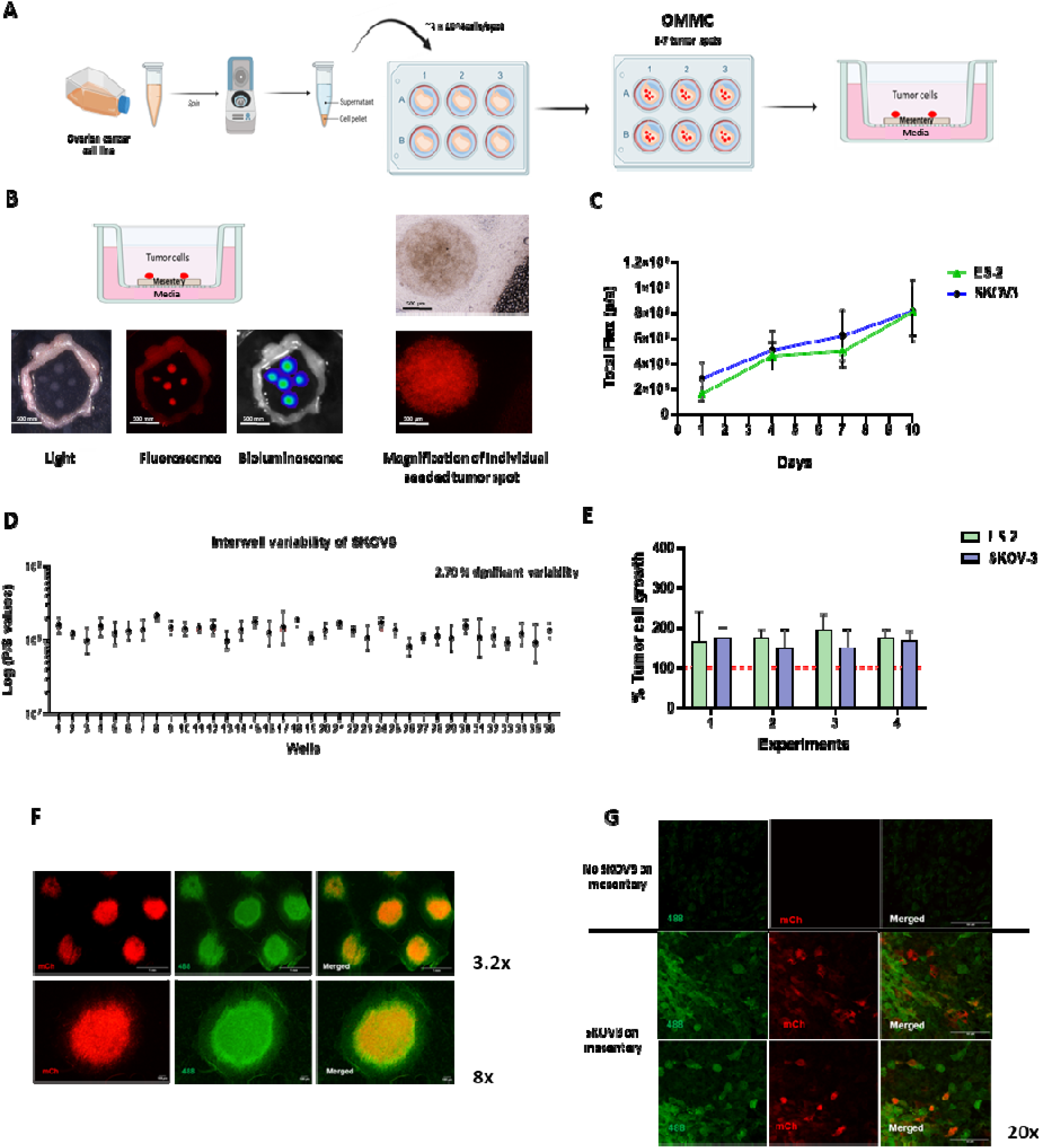
Tumor seeding on OMMC, growth, and tumor-mesentery interaction. (A) Process of tumor spot seeding on OMMC. (B) Light, Fluorescent, and BLI pictures from above view of Day 3 seeded tumor spots on a mesentery with a magnified display of a well-rounded tumor spot. (C) ES-2 and SKOV3 showed consistent tumor growth on OMMC over 10 days. (D) Minimal inter-well variability after manual tumor seeding on Day 0 (>600 multiple comparisons for 36 wells). (E) Consistent and reproducible tumor growth of ES- and SKOV3 across separate experiments measured on Day 5 showing survival and proliferation. (F) Representative fluorescent images (3.2 x and 8x) of seeded SKOV3 (Day 3) tumor spots (red-mCherry), CD11B (green-Alexa Fluor 488) and merged. (G) 20X magnification on the tumor spot edges on the mesentery (includes mesentery with no tumor, first row) showing a clear macrophage activation when the tumor is present.

The mesentery and omentum are known to contain blood vessels, lymphatic vessels, resident macrophages, and fibroblasts all embedded within extracellular material such as collagen and elastin fibers (*27,28*). We next investigated tumor-mesentery interaction by exploring the potential activation of macrophages residing in the mesenteric tissue in the presence of OC foci. When mesenteric tissue with OC foci was stained 72 hrs post-seeding with anti-CD11b antibodies, immunofluorescent imaging revealed a robust signal co-localized with the OC foci (Fig. 3F-G), an effect that was not observed in mesentery without OC, suggesting that the presence of OC tumors induced recruitment and activation of macrophages around the tumor foci. High-magnification images of this interaction revealed that CD11b-positive cells were indeed a distinct population from mCherry tumor cells, and that a “halo” of activated macrophages existed a short distance wider than the tumor cell footprint on OMMCs.

### Developing OMMCs as a drug-testing platform

Building towards the development of a new drug testing platform, we next investigated drug responses within the OMMC system. Using the PI assay, we first measured the off-target toxicity of a panel of therapeutics by exposing OMMCs to a panel of FDA-approved agents for the treatment of OC. The panel included olaparib, gemcitabine, carboplatin (carbo), paclitaxel (pac) and carboplatin-paclitaxel (carbo-pac). We dosed at increasing drug concentrations of 0 - 1000uM as single agent therapies or using combination regimens (full dose range of carbo-pac at 10uM [carbo-pac10] or 100uM [carbo-pac100]). Our PI assay showed the single-agent treatment induced no significant decreases in the viability of the mesenteric tissue. The combination regimen also showed similarly low toxicity at lower doses, but reductions up to 25% in mesenteric viability were detected at higher dose ranges and showed higher variability (Fig 4A).

**Figure 4.**
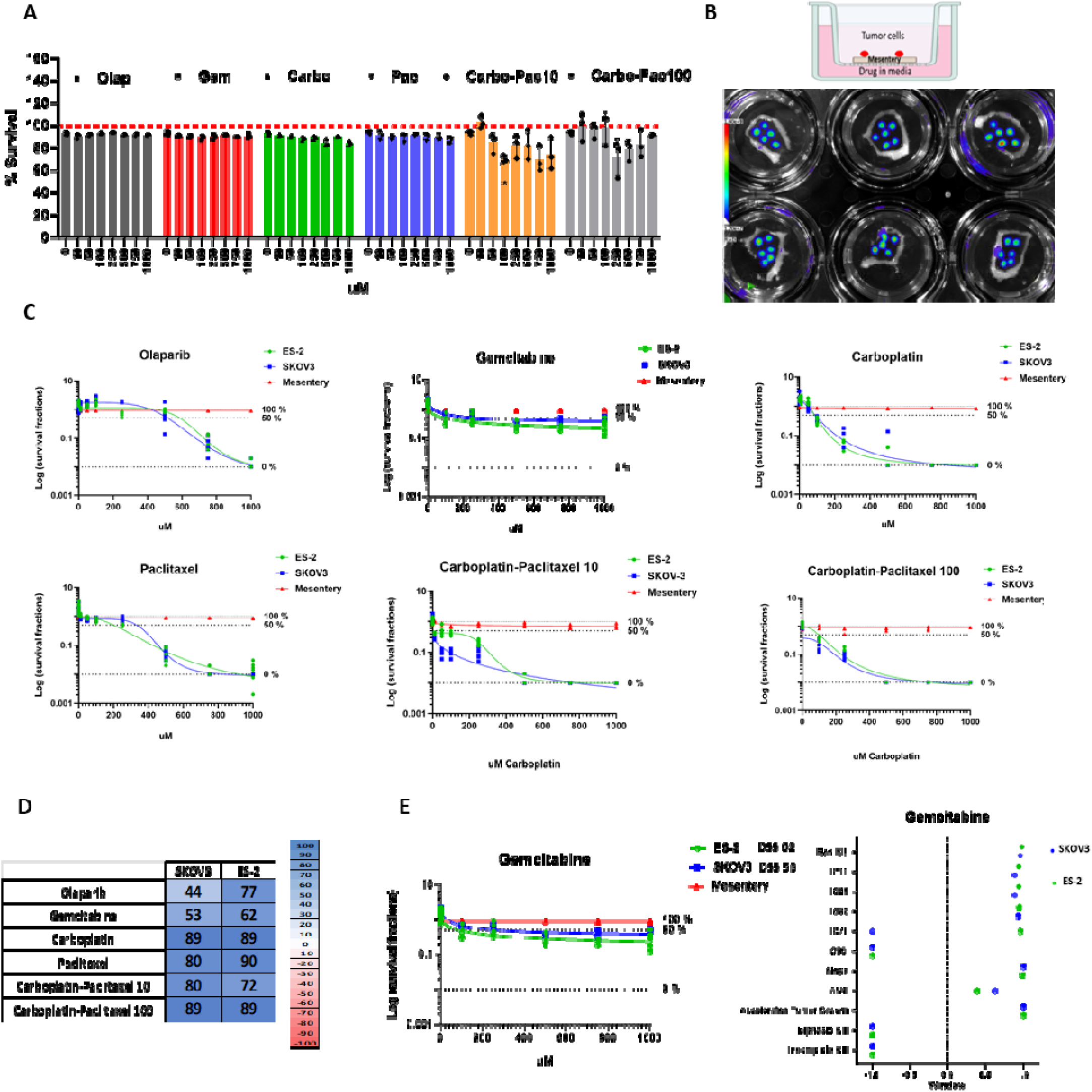
Tumor drug response, toxicity on the mesentery, and drug sensitivity score (DSS). (A) Drug effect on survival of tumorless mesenteries, when exposed to increasing concentrations of FDA-approved single and combination chemotherapies for 3 days. One-way ANOVA p>0.05, p=0.032 (B) Visual of schematic (Top) and real (Bottom) OMMC system with ES-2 tumor spots suggesting its potential functionality to assess tumor drug response by BLI quantification. (C) Tumor drug-response curves and mesentery viability after 3-day exposure to the same group of chemotherapies. (D) Calculated DSS for ES-2 and SKOV3 against all drugs. 0 to 100 suggest increasing efficacy in tumor kill relative to OMMC toxicity, while scores from 0 to −100 describe scenarios in which tumors thrive more effectively than OMMC for a given treatment (E) Example of the tumor response curves for the two cell lines against Gemcitabine and the corresponding DSS. (F) Example of the therapeutic window across all DSS weighted parameters for Gemcitabine on the treated cell lines. Values ranged from −1 to +1, where values approaching +1 indicate better tumor kill relative to less toxicity on the tissue, and values approaching −1 suggest tumors remained viable while toxicity to the normal OMMCs tissue was elevated. See supplemental figure S4 for each treatment.

Next, we focused on the tumor response and seeded the ES-2 and SKOV3 cell lines directly atop the mesenteric tissue. We assessed drug activity against the OC foci by quantitative bioluminescence imaging (BLI) (Fig. 4B). Dose-response curves and half-maximal inhibitory concentration (IC_50_) values were then calculated for each therapeutic agent and the best-fit dose-response curves for ES-2 and SKOV3 killing and off-target OMMC toxicity were generated (Fig. 4C). See also supplemental figures for individual tumor response curves per cell line (Fig. 4SA)

We then utilized our previously developed normalized scoring system to calculate “Drug Sensitivity Scores” (DSSs) for each drug-tumor-OMMC interaction (*25*). This summarized scoring system compares the dose-response curves for tumor kill and OMMC toxicity for each therapeutic to calculate “therapeutic windows” across 11 different characteristics of curves such as the IC_50_ and the Area Under the Curve (AUC). The therapeutic window ratio of tumor kill to OMMC toxicity for each parameter can range from −1 (low tumor kill but high OMMC toxicity) to +1 (high tumor kill and low OMMC toxicity), and the final DSS value is calculated from a weighted, multipoint mathematical algorithm that collapses all 11 parameters into a single score. DSS values can range from −100 to +100, where +100 describes the maximal performing agents (greatest tumor kill, lowest toxicity) and −100 describes the lowest performing agents (poorest tumor kill, highest toxicity). These unique values represent an overall metric that captures the drug response across tumor foci detected by BLI and the changes in the viability of normal mesentery tissue measured by our predefined PI assay, allowing drugs with different potencies to be compared with less bias toward more potent compounds.

OC cell lines ES-2 and SKOV3 were similarly sensitive to single-agent therapy with carbo and the combination carbo-pac 100 and thus recorded similar DSSs; however, variable DSS values were calculated for the combo carbo-pac 10 and single-agent therapy with gemcitabine, pac, and olaparib. Focusing on the Gem treatment as an example, a DSS of 53 for SKOV3 and a DSS of 62 for ES2 (Fig. 4D) were calculated, indicating that a slightly better overall tumor-killing effect on ES-2 took place. This result was driven by differences in AUC between the two tumor response curves and the inability of Gem to kill at least 75% of SKOV3 cells (Fig 4E). (See also Fig. S4B).

### Advancing the OMMC platform to incorporate fresh OC patient tissue

A patient’s own live tumor tissue best represents their own disease, but models which functionally assess drugs using uncultured tumor tissue have proven challenging. Often, these models require a long lead time and conditions that impact phenotypic changes, marked clonal selection, and genetic drift that results in loss of the cellular and genetic heterogeneity present in clinical OC patient tissue. To develop OMMCs into a model that fulfills these needs, we designed a method to prepare and engraft a diverse panel of living, uncultured patient OC tumor tissues onto OMMCs for rapid, functional drug screening and eventual treatment guidance. Toward that end, we obtained fresh surgically resected OC tumor tissues from patients undergoing standard-of-care resection surgeries at the University of North Carolina at Chapel Hill (UNC-CH) hospitals following informed consent on the UNC-CH MASCOT trial (LCCC 1855 MASCOT: Manufacturing and Analysis of Stem Cells from Skin Cells for Ovarian Cancer Treatment). After cryopreservation and thaw of the tumor tissue according to our optimized protocol, the tissue was mechanically dissociated into a homogeneous near single-cell suspension, rapidly transduced with LV-mCh-FLuc, and seeded as concentrated tumor foci, with each containing a representative sample of ∼0.5 mg tissue onto OMMCs.

To measure the ability of OMMCs to engraft patient OC tumor tissue, we explore tumor survival and growth through normalized BLI signals from Day 6 to Day 3 of tumor samples from nine individuals Fig. 5A). Results showed all 9 tumor tissues remained viable with mean values of fold tumor growth above one during this time. We also compared 6 of these tumor tissue engraftments (1) on OMMCs, (2) in transwell culture without OMMCs, and (3) in standard *in vitro* culture via BLI (Fig. 5B and 5C). Culturing the tissues across different formats revealed that the same tissue cultured *in vitro* or *ex vivo* without mesentery exhibited minimal or undetectable viability when compared to our OMMC system where they significantly thrived. We then investigated the reproducibility of the tumor spot placement per well at Day 0 using one OC patient tissue (Mascot Patient # 62). After performing multiple inter-well comparisons across 30 mesenteric tissues seeded with 5 tumor foci each, results showed high consistency, with only a 3.68% difference across tissues with tumors (Fig. 5D)

**Figure 5.**
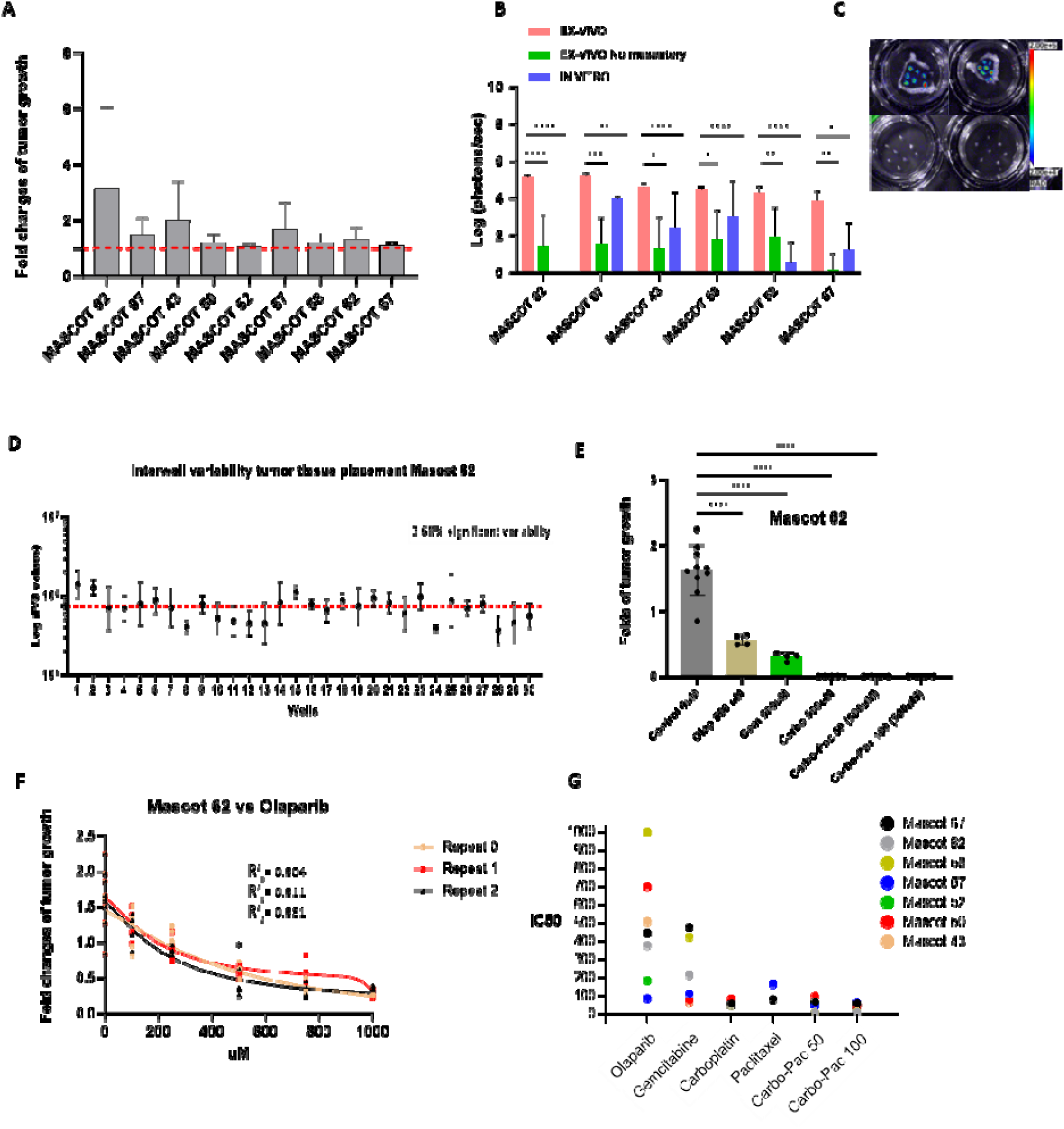
Human ovarian tumor tissues on OMMCs and tumor response. (A) Mean values of the tumor growth fold on OMMCs, showing all patient tumors stay alive and even proliferate for some in 6 days. (B) Comparison of patient OC tumor growth in different cultured systems. Two-way ANOVA, ****p < 0.0001, *** p < 0.001, **p < 0.01, *p <0.05. (C) Representative BLI picture of patient tumor spots when mesentery is present or not, where clear tumor viability is observed. (D) Example of inter-well variability of the human OC tumor spots at the time of placement on the mesentery membrane, showing there was a consistent tumor cell manipulation with no significant difference in BLI values inter-well. (E) Significant tumor response on OMMCs to 500uM of chemotherapies. One-way ANOVA ****p<0.0001 (F) Example of reproducibility on the Mascot 62 tumor tissue response to Olaparib exposure for 3 days. One-way ANOVA, *p <0.05 (G) Graph representing different drug potencies at IC_50_ for each of the patient tumor tissues engrafted on OMMC.

We next tested the responses of patient tumor tissues when exposed to 500uM of olaparib, gemcitabine, carbo, and carbo-pac 50 and 100. Results showed clear differences in the potency of tumor responses for each patient (Fig. 5E and Fig. S5A). Also, consistent reproducibility resulted when a series of experimental repeats were performed, using a full therapeutic dose range (0 to 1000 uM) of olaparib for one of the patients (Fig. 5F). All drugs used in this study were then tested against each patient tumor tissue (Fig. S5B), and IC_50_s for each treatment among all tumors were plotted (Fig. 5G), with lower IC_50_s indicating greater tumor sensitivity.

Although IC_50_s give an informative measure of the efficacy of drugs, it is not the only parameter analyzed to account for drug sensitivity in our OMMC platform. We grouped all the patient tumor response curves by treatment and therapeutic window ratios to better visualize drug efficacy among patients and parameters (Fig. 6A). We then calculated the DSS values following the same multi-parametric algorithms described previously, collapsing all normalized parameters into a single score simplifying the comparison of each drug-tumor-OMMC interaction (Fig. 6B).

**Figure 6.**
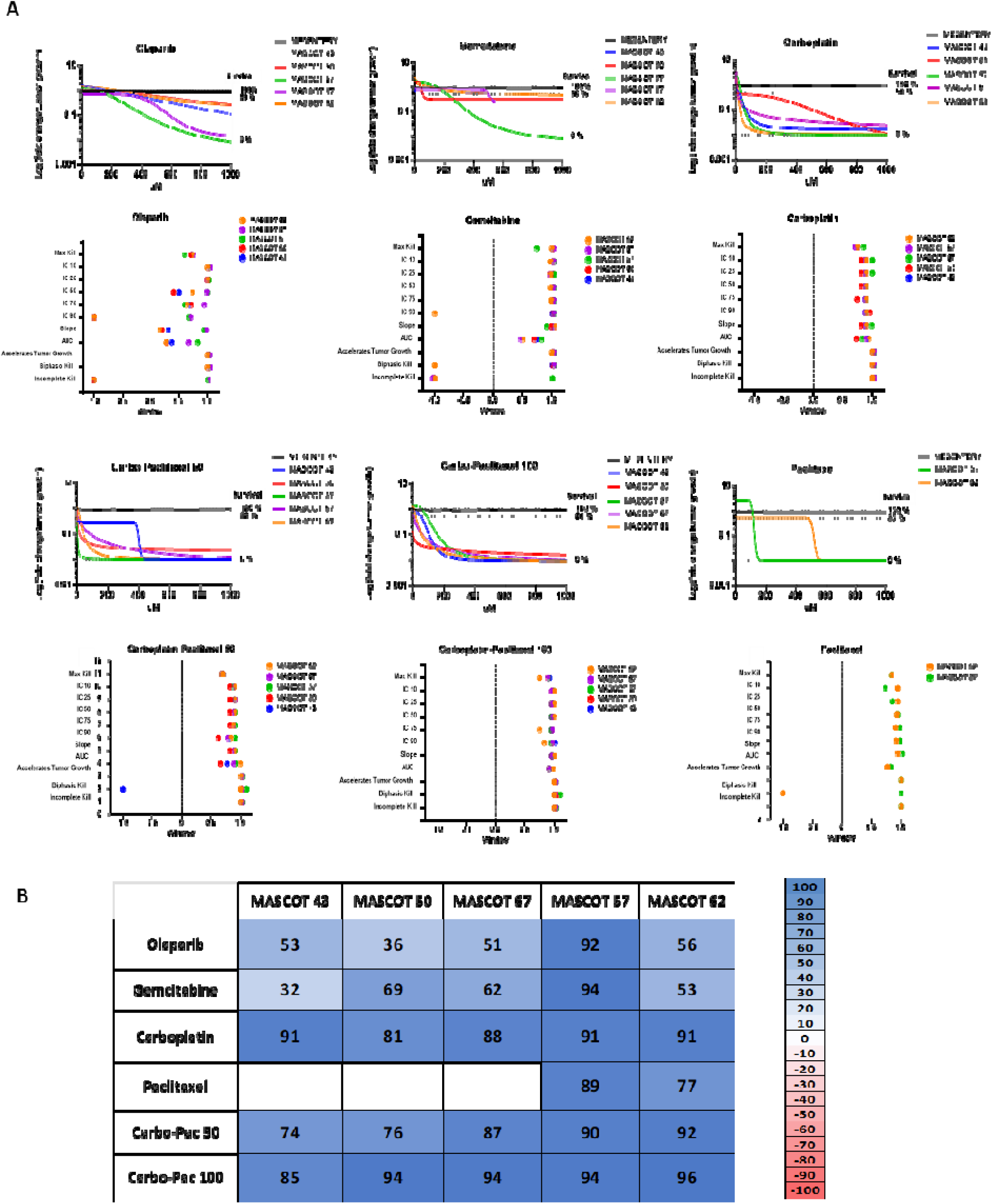
Patient tumor tissue responses to all treatments and calculated DSS. (A) Tumor response curves on OMMCs per individual treatment (top) and their corresponding therapeutic window across all parameters (bottom). (B) DSSs array for both cell lines against alt drugs. Values from 0 to 100 suggest increasing efficacy in tumor kill relative to OMMC toxicity, while scores from 0 to −100 describe scenarios in which tumors thrive more effectively than OMMC for a given treatment.

### OMMC-derived patient sensitivities and clinical data

The above data suggests that our OMMC functional precision medicine platform is a promising strategy to guide clinical decisions. Here, we begin to associate the treatment responses predicted by our OMMC assay and the actual clinical response from five OC patients. We, therefore, collected data on tumor type, genetic mutations, treatments received, tumor recurrence, etc., of each patient and compared to DSSs of their tumor tissues after treatment on OMMCs (Fig 7). In the clinic, the OC patient’s tumor responses to drug therapy are confirmed by the presence of clinical symptoms, imaging and blood levels of CA125. In our OMMC, the calculated DSSs tell us how the patient’s tumor responds to each treatment and allow us to establish relative comparisons between these values from different treatments within one patient and between different patients with the same treatment. In this study, we collected all individual tumor biopsies on the day of surgery, later the processed tumors were seeded on the OMMC and exposed to the single therapies olaparib, gemcitabine, and carbo, in addition to the combination carbo-pac. To later discuss potential associations between results from the clinical and the OMMC, we decided to each case separately.

**Figure 7.**
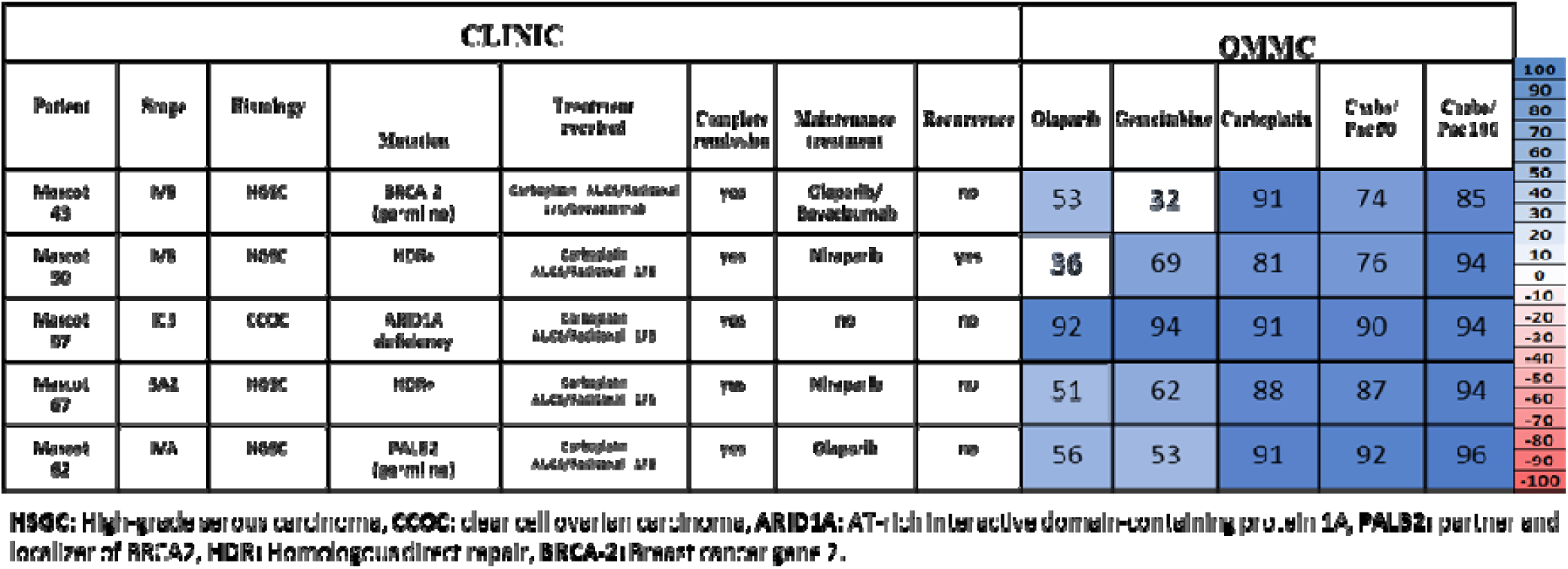
Clinical data and OMMC results. For the calculated DSSs, from 0 to 100 signify increasing efficacy in tumor kill relative to OMMC toxicity, while scores from 0 to −100 describe scenarios in which tumors thrive more effectively than OMMCs for a given treatment

MASCOT Patient #43’s tumor was identified as IVB stage, HSGC with BRCA2 germline type mutation, and received six cycles of neoadjuvant chemotherapy (NACT) carbo-pac before surgery to which partially responded later showing complete remission after the frontline treatment (NACT + surgery). On OMMC, the calculated DSSs showed a relatively stronger tumor response to carbo (DSS:95) and carbo-pac 100 (DSS:85) than to the rest of the treatments: olap (DSS: 53), gem (DSS:32) and carbo-pac 50 (DSS:74). In clinic, after surgery, this patient responded well to 6 cycles of adjuvant chemotherapy (ACT) of carbo-pac followed by maintenance therapy regime with olaparib and bevacizumab with no tumor recurrence within 6 months.

MASCOT Patient #50 tumor was identified as IVB stage, HSGC as well but with HDR+ mutation and received six cycles of NACT carbo-pac before surgery. Partial response to this treatment was observed and a complete remission to the frontline treatment was recorded. From our platform, calculated DSSs showed also a relatively stronger tumor response to carbo (DSS:81) and carbo-pac 100 (DSS: 94) to the rest of the treatments: olap (DSS: 36), gem (DSS:69) and carbo-pac 10 (DSS:76). This patient received 6 cycles of carbo-pac after frontline treatment and maintenance therapy of niraparib after the ACT was complete, however, did recur with brain metastasis within four months of treatment with this PARP inhibitor.

MASCOT Patient #57’s tumor was classified as CCOC at stage IC3 and there was no mutation in BRCA or HDR genes detected but showed ARID1A deficiency on next-generation sequencing standard-of-care tumor testing. This patient was not treated with NACT before surgery however a complete remission was recorded. It is noteworthy that at the time of tissue collection, this patient’s tumor was chemo-naïve when being tested on the OMMC and showed a high tumor response to all treatments as the calculated DSSs reflected in Figure 7. This patient received 6 cycles of carb-pac after surgery but not maintenance therapy at all and did not recur after 6 months of treatment.

MASCOT Patient #62’s tumor was identified as IVA stage, HSGC with PLB2 germline mutation, and received four cycles of NACT before surgery. Clinical partial response to this treatment was observed and a complete remission to the frontline treatment was recorded. On the OMMC this patient’s tumor had relatively higher response to carbo (DSS:91), carbo-pac 10 (DSS:92) and carbo-pac 100 (DSS: 96) to the rest of the treatments: olap (DSS: 56), gem (DSS:53). In clinic this patient underwent 6 cycles of carbo-pac after frontline treatment followed by a maintenance therapy of olaparib. This patient did not recur within six months of treatment with PARP inhibitor as maintenance therapy.

MASCOT Patient #67’s tumor was identified as 3A2 stage, HSGC with HDR+ mutation, and did not receive the NACT of carbo-pac before surgery however showed a complete remission. On the OMMC this patient’s tumor being chemo-naïve was exposed for the first time to drug therapy. Calculated DSSs reflected a higher response to carbo (DSS:88), carbo-pac 10 (DSS:87) and carbo-pac 100 (DSS: 94) than the rest of the treatments: olap (DSS: 51) and gem (DSS:62). In clinic this patient underwent 6 cycles ACT of carbo-pac after surgery followed by maintenance therapy of niraparib and did not experience tumor recurrence within six months of treatment.

## Discussion

Although OC treatment decisions are mainly based on histologic classifications, in combination with specific tumor genetic mutations, there is a clear percentage of patients who do not show a clinical response to standard treatments, despite having histology and molecular alterations that are biomarkers for response (i.e., high grade serous and high response to carbo-pac or BRCA mutations and PARP inhibitors) (*29–35*). Our studies describe a new tool to *functionally* predict how each tumor will respond to therapy and maximize the survival of patients with OC.

After a meticulous optimization process during OMMC generation, we showed the selected region of interest on the rat mesentery is consistently viable up to 10-11 days independently of animal age and across separated experiments. Reproducibility experiments showed established human OC cell lines survived and proliferated on the mesentery membrane during the time studied (10 days), while human tumor tissues thrived in our OMMC, in contrast to *in vitro or ex vivo* (no mesentery) conditions. This coupled with a marked tumor spot-mesentery interaction observed by the activation of immune cells, might suggest a microenvironment of a living substrate is necessary to establish certain biological conditions for optimal tumor tissue engraftment and survival. Therefore, we believe our platform facilitates the study of these difficult-growing resected human tumor tissues in the laboratory and demonstrates the existence of a dynamic and responsive living system such that an intact tumor and tissue substrate microenvironment is preserved.

Our OMMC system allowed us to test tumor responses of established human OC cell lines and tumor tissues surgically dissected from patients to FDA-approved standard-of-care treatments for OC patients. Consistent reproducibility on tumor response was proven showing the functional reliability of our OMMC. Simultaneously, because we could also measure the impact of drug toxicity on the healthy mesenteries, we were able to calculate DSS for each treatment within each cell line or tumor tissue, in which a series of 11 kinetic parameters were taken into account to reflect a more complete measure of the treatment efficacy and potentially facilitate a tool to guide precision medicine in the future. The DSS values per cell line and treatment suggested similarities in tumor kill effectivity for single carbo and its combination (carbo-pac 100) (Fig. 4D). Lower and variable DSS values from Gemcitabine treatment were seen when compared with taxane-platin-based therapies (ES-2 DSS: 62, SKOV3 DSS 53). In addition, ES-2 was more sensitive to the PARPi olaparib than SKOV3 (ES-2:77, SKOV3:44). These results from our OMMC are aligned with *in vitro* data generated by the Genomic of Drug Sensitivity in Cancer Project (*36*). These two cell lines are classified as different tumor types: SKOV3 is a non-high-grade serous carcinoma cell line and ES-2 is an unusual clear cell adenocarcinoma that appears more like high-grade serous carcinoma (*37*). However, there are some similarities in the genetic profiles of these two well-known OC cell lines such as both being TP53 mutant, BRCA wildtype, and HRD negative. Given this, we might have expected both cell lines to exhibit less sensitivity to PARPi treatment. Thus, these results suggest that we should not just consider *BRCA*- and *TP53*-mutations or HRD status (*38*) to delineate the sensitivity of different drugs to PARPi and other chemotherapeutics, but the entire mutational profile of the potential oncogenes for each cell line.

In our small cohort study with 5 patients, the OMMC allowed us to elucidate preliminary associations between the ex vivo patient’s tumor responses reflected in the DSSs and the results from the clinic after being exposed to similar treatments. Since we collected tumor biopsies on the day of surgery, some patients had already received NACT of carbo-pac, but not additional chemo or maintenance therapy, therefore this suggests residual tumors after this NACT were tested on the OMMC. To better discus potential associations, we focused on 1) the clinical outcomes (complete remission or not) of each patient from the NACT and their tumor responses (DSSs) on the OMMC and 2) the clinical outcomes postsurgery (recurrence or not) after the ACT (carbo-pac) followed maintenance therapy (PARPi and/or bevacizumab) and the DSSs of these treatments revealed from OMMC.

1) Mascot Patients #43, 50, and 62 went into remission after frontline treatment (NACT of carbo-pac and surgery). In our OMMC these tumors responded well to carbo and carbo-pac treatment when we established relative comparisons of their DSSs to the other treatments (rows fig. 7) and also showed homogenous tumor responses among patients (columns fig. 7). These results aligned with the patient’s clinical outcomes, showing a positive association between these two separate results. It is well-known that the majority of OC patients (up to 85%) can achieve remission after front-line carbo-pac chemotherapy in combination with cytoreductive surgery, regardless of whether chemotherapy is given in a neoadjuvant or adjuvant manner. MASCOT Patients #57 and 67, also went into complete remission but they only underwent debulking surgery at that point without NACT; however, their tumors were treated on OMMC with taxane-platin based as the rest, and similar strong responses were observed.

In the OMMC, we included gemcitabine, as a nucleoside metabolic inhibitor, a common second-line agent for OC treatment. Since none of these patients was treated with gem in the clinic we could not explore associations with our OMMC but Interestingly, if we focused on the DSSs of gem, carbo, and carbo-pac of all patients, our ex vivo platform found heterogeneity in tumor responses to gem among different patients (DSSs columns fig. 7), in contrast to relatively homogeneous responses from carbo-pac-based therapies. This aligned with the usual greater sensitivity of OC to frontline carbo-pac treatment as opposed to second-line gemcitabine therapy. This evidences that the OMMC’s functionality can detect these differences being in synchrony with the documented findings in the clinic.

2) After cytoreductive surgery, for patients with biomarkers for response to PARPi (either somatic or germline BRCA mutations or HRD), maintenance therapy with PARPi +/- bevacizumab is recommended after 6 cycles of ACT. In our OMMC we included olaparib as the only PARP inhibitor however in clinic either olaparib or niraparib was used. If we compare the DSSs of olaparib with the DSSs of platin-based treatment (OMMC section, rows 1-6 Fig. 7) of the same patient and for all patients it is noted they were lower for the four patients classified as HGSC (Mascot Patients #43,50,62 and 67) suggesting less relative efficacy than carbo and its combination carbo-pac. Then, if we look closely at the DSS values for olap across patients (OMMC section, column 1 Fig. 7) 53, 38, 51, and 56 for Mascot Patients #43,50,67 and 62 respectively, we noticed Mascot Patient #50 had the lowest (DSS of 38) while the other ones were similar. In the clinic, Mascot Patient #50, had a tumor with HRD that should be predictive of response to the PARPi treatment, however, this patient did recur within four months. These findings might confirm the importance of the functional drug testing capabilities of OMMC since the clinic outcome suggests the maintenance therapy with olap was not enough to avoid recurrence while our results from the OMMC reflected the lowest DSS of this treatment among all patients. This might also suggest that our OMMC platform could potentially provide insights on tumor recurrence prediction although further studies are needed.

The tumor from Mascot Patient#57 was classified as a stage 1C, CCOC. In the OMMC, the tumor tissue of this patient showed a strong response to all drug therapies used and the highest DSS values for olaparib and gemcitabine among the rest of the patients, meaning potentially greater sensitivities for these drugs. In clinic this patient did not recur. Studies have reported CCOC tumors can be sensitive to PARP inhibitors even in the absence of BRCA/HDR mutations (*39*), however, CCOC are known to be resistant to platinum-based chemotherapy, while a small subset shows a positive response (*40,41*). This patient tumor tissue responded well to carbo and carbo-pac based therapy in the clinic and our platform. CCOC is characterized by genetic alterations distinct from those found in HGSC and the therapeutic options with PARPi are limited due to the low frequency of BRCA1/BRCA2 mutations in these cancers (*42*). Consequently, a great deal of attention in OC cell type has been paid to synthetic lethal therapies that target vulnerabilities conferred by ARID1A deficiency. Like the BRCA1 and BRCA2 proteins, BAF250A/ARID1A promotes homologous recombination-mediated repair of DNA double-strand breaks, suggesting that PARP inhibition might be therapeutically effective (*43*). Clinical trials of the PARP inhibitors olaparib and niraparib, using ARID1A deficiency as a biomarker, are underway in ovarian and other cancers (NCT04065269, NCT04042831, NCT03207347). Mascot Patient #57 did have had a tumor ARID1A deficiency which may explain sensitivity to PARPi treatment on the OMMC platform. Here again, we have demonstrated that results from OMMC aligned with the clinical outcomes in this case as well as suggesting our OMMC could be a potent tool to carry out functional drug testing for precision oncology.

While we continue to work on the next optimization level for our OMMC and increase our clinical study population we believe that a need to add functional testing to the already existing clinical diagnostic landscape is fundamental since we might predict with higher accuracy a more appropriate and personalized therapy for OC patients. While debulking surgery and all these chemotherapy regimens with neoadjuvant, adjuvant, and maintenance therapies may show strong and rapid responses in some patients we should not assume a similar response type for other patients which can differ from person to person and may also lead to different outcomes. Currently, specific genetic mutations, disease stages, tumor type, overall health condition of the patient, tumor heterogeneity, optimal or suboptimal debulking surgery, and age are the main factors to consider when deciding the right treatment. Here we show a novel ex-vivo platform for OC with the potential to functionally test a variety of treatments that ultimately would help to guide medical treatment decisions. We believe the preliminary associations between the results observed in the clinic and our results coming from our OMMC support the functionality of our platform and the need for further exploration of this novel tool in OC management.

## 2. Materials and methods

### Animals

Adult dams and 3-, 4-, 8- and 16-week-old female Sprague-Dawley rats from Charles River Laboratories were used in this study. The animals were housed in sterile rooms of an accredited AAALAC laboratory animal facility at the University of North Carolina-Chapel Hill (UNC-CH) and subjected to procedures described in the correspondent animal protocol and approved by its Institutional Animal Care and Use Committee (IACUC).

### Surgical procedure and mesentery collection

All surgeries performed were non-survival. The rats were anesthetized and euthanized using an overdose of 5% isoflurane for a period of 10 minutes followed by a second method of euthanasia: cervical dislocation on rats weighing less than 250g and thoracotomy on dams. The rats were placed in a prone position and the skin on the abdominal area was disinfected with 7.5% iodine followed by 70% ethyl alcohol. A longitudinal incision of the skin and the abdominal muscle was made along the midline starting right above the urinary aperture up to the xiphoidal process of the sternum in the thoracic region. Subsequently, the intestines were exposed, and the mesentery’s regions of interest were identified and dissected using sterile micro scissors and forceps. 10 to 12 mesenteries membranes were removed aseptically from each rat. These isolated mesentery tissues were at once placed in culture media as subsequently described.

### Organotypic Mesentery Membrane Culture System (OMMCS)

0.4µm porous membrane cell culture inserts (Millicell Cell Culture Insert, 12 mm diameter, hydrophilic PTFE, 0.4 µm pore size from MilliporeSigma) were placed into wells of 6-well plates containing 1 mL of minimum essential medium (MEM, Gibco^TM^) supplemented with 10% fetal bovine serum (FBS) (Avantor®Seradigm, Premium Grade) and 1% penicillin-streptomycin (P/S) (Gibco^TM^). Each dissected mesenteric tissue was placed on a cell culture insert in contact with the culture media. With the use of forceps, mesenteric tissues were spread on the surface of each insert without touching the mesenteric membrane. All 6-well plates with mesenteric tissues were then transferred to an incubator under standard conditions of 37℃,5% CO2, and 95% humidity.

### Mesentery viability measured by Bioluminescence

Six freshly prepared mesenteric membranes with an approximate diameter of 1.5cm were incubated with 1µl polybrene and 2ml of a lentivirus with the mCherry fluorescence reporter protein fused to firefly luciferase (LV–mCh-FL) at 7.5e6 vg/ml at 37°C for 24h. After incubation, the mesenteric tissue was washed thrice with 1X Dulbecco’s phosphate-buffered saline (PBS) to remove the residual virus. The mesenteric tissue was then replated in fresh MEM culture media supplemented with 10% FBS (Avantor®Seradigm, Premium Grade)1% P/S (Gibco^TM^). On days 4, 6, 8, and 10 after initial mesenteric membrane plating, the bioluminescence intensity of each transduced membrane was measured by the Ami HTX imaging optical system (Spectral Instruments Imaging) following 5 minutes of exposing the tissue to 1ml at 0.375mg/mL high-quality IVISbrite^™^ D-luciferin Potassium Salt (PerkinElmer^TM^) substrate mixed in the culture media.

### Mesentery viability and tissue health condition measured by Fluorescence using Propidium Iodide assay (PIA)

By measuring the fluorescence emitted by dead cells in the mesentery, PIA was used to assess the tissue health condition across experiments, the viability over time of fresh tissue isolated right after dissection, and the potential local toxicity of each treatment the membrane was exposed to. Mesentery viability of various rat ages (3, 4, 8, and 16 weeks old) and dams were compared to select the optimal animal age to be used for the OMMCs in this study. Toxicity was assessed at the end of each therapy assay: t=3 days after the initiation of each treatment and t = 5 after OMMC generation. Propidium iodide (PI) (Sigma-Aldrich ®) was added to the media in each well for a final concentration of 5ug/mL and after an incubation period of 1 h, the fluorescence intensity was determined using the Ami HTX imaging system (Spectral Instruments Imaging). Positive control was generated by killing the OMMCs via incubation with 100% DMSO (Dimethyl sulfoxide) for 3 hours. A killing curve was generated after the tissue was exposed to increasing concentrations of DMSO (0.5, 1, 3, 5, 10, 20, 50, 75, and 100%) in the culture media.

### Mesentery cell count per region, and membrane thickness

5 mesenteries of 8-week-old rats with an approximate surface area of 1.76cm^2^ were selected and 5 regions in each mesenteric sample were randomly studied to estimate the thickness and the number of cells in the membranes belonging to the Jejunum and Ileum separately. A total of 10 mesenteries were killed by exposition of 100% DMSO for 3 days and PIA was applied to stain all dead cells. A confocal laser-scanning microscope (Zeiss 780) from UNC-CH Neuroscience Microscopy Core was used to measure the thickness and number of cells of the mesenteric membrane. A Z-stack for the thickness was defined as a series of images captured at incremental (4μm) steps in the Z plane.

### Cell lines and culture

Human ovarian cancer well-established cell lines SKOV-3 (non-high grade serous carcinoma cell line) and ES-2 (unusual clear cell adenocarcinoma but appear more like high-grade serous carcinoma) (*64*) were obtained from the Lineberger Cancer Center Tissue Culture Facility of the University of North Carolina. Transduction of these cell lines with lentiviral vectors encoding optical reporters mCherry-firefly-luciferase (LV-mCherry-FLuc) was carried out in our laboratory. SKOV-3 and ES-2 cell lines were cultured and propagated in McCoy’s 5A media (Gibco^TM^) supplemented with 10% FBS (Avantor®Seradigm, Premium Grade) and 1% P/S (Gibco^TM^) at 37℃ in a humidified atmosphere of 5% CO2.(2,3). The cells were used at 80% to 90% confluency for all the experiments with cell viability ranging from 95 to 98% (Countess II FL, Invitrogen, Thermo Fisher Scientific).

### Ovarian cancer cell lines on OMMCs

SKOV-3 mCherry-Fluc and ES-2 mCherry-Fluc cells were seeded onto OMMCs in the form of tumor spots, 2h after tissue collection. Each mesentery in between 5 and 7 tumor spots consisting of a suspension of ∼2.0 x10 ^4^ cells in 1uL. To measure tumor cell viability, D-Luciferin was added underneath the transwell insert and allowed to incubate for 5 minutes before bioluminescence (BLI) measurement was taken on an AMI optical imaging system at days 4,6,8 and10 for the survival experiments and at days 2 and 5 for the tumor drug response studies.

### Immunohistochemistry (IHC)

IHC was performed using the primary antibody for a cluster of differentiation molecule 11B (CD11B [Abcam, ab133357], 1:250. Day 1 cultured mesenteries without tumor spots and Day 5 cultured mesenteries with SKOV-3 mCherry-FLuc tumor spots were fixed in the same culture plates, with 4% paraformaldehyde, and stored at 4°C. After 48 hours of fixation, the mesenteric membranes were carefully washed 3 times with PBS and then stored at 4°C until IHC was conducted. Samples were then washed for 10 minutes in 0.1% triton X-100 in PBS at room temperature, then blocked in 5% fetal bovine serum (Avantor®Seradigm, Premium Grade) in PBS for 1h at room temperature. The samples were incubated for 24hrs at room temperature in a primary antibody solution (CD11B) and a blocking buffer with a rotating motion then washed 3 times in PBS for 10 minutes. Next, the mesenteries were incubated with a secondary antibody consisting of a blocking buffer solution and Alexa Fluor 488 goat anti-rabbit IgG (Thermo Fisher Scientific, A-11008, 1:1000) for 1 hour in a darkroom then washed 3 times with PBS for 10 minutes and mounted on microscopic slides. Liquid mounting (ProLong^TM^ Gold Antifade Mountant, Cat: 10144) was applied, coverslips were added, and slides were allowed to cure overnight. Z-stack images were acquired using a Zeiss 780 confocal microscope at UNC-CH Neuroscience Microscopy Core. Z-stack images are processed by converting them into maximum intensity projection (Max IP) images. The brightness of Max IP images was then further adjusted to accurately assess and present the morphological changes of macrophages.

### Ovarian cancer cell lines drug response on OMMCs

Two days after initial tumor cell seeding on OMMCs, an initial BLI reading (Day 2) before treatment, was performed as described above. All the wells and transwell inserts in the culture plate were washed 3 times with PBS right after the BLI was done to remove the remaining luciferin added initially. The transwell inserts were carefully rinsed from the outsides with PBS without disturbing the position and integrity of the mesentery tissue holding the tumor spots. Fresh MEM 10% FBS and 1% P/S culture media were added underneath each transwell insert after the washes. Next, six concentrations (0,10,50,100,250,500,750 and1000 µM) of each drug were diluted in the fresh media (n = 5-7 tumor foci per concentration; n=40 to 56 tumor foci per drug per cell line). The plates were put back in the incubator at 37℃ in a humidified atmosphere of 5% CO_2_ for three days when a second BLI reading (DAY 5) was performed. Dose-tumor response curves were generated. Data values were normalized to DAY 2 values for all groups and transformed into survival percentages. Each group was compared to an untreated control group.

### Chemotherapies

Paclitaxel from Selleck Chemicals (S1150|CAS: 159634-47-6), olaparib from Selleck Chemicals (S1060|CAS: 890090-75-2), and gemcitabine from ApexBio (A8437|CAS: 95058-81-4) were reconstituted in dimethyl sulfoxide (DMSO, Fisher BioReagents) in an optimal percentage range (0.5 to 3 %). carboplatin from Sigma-Aldrich (C2538-100MG|CAS: 41575-94-4), was reconstituted in PBS only and used immediately after. At day 2 of ovarian cancer cell incubation on OMMCs, monotherapies with carboplatin, paclitaxel, olaparib, gemcitabine, and combination therapies with carboplatin-paclitaxel were carried out for 3 days. For single therapies, drugs were added in the culture media below the transwell inserts at final concentrations of 10, 50, 100, 250, 500, 750 and 1000 µM. For combinations, the same dose range for carbo in single therapy but with paclitaxel (10, 100 µM). Bioluminescence images were taken on an Ami HTX imaging system as described above.

### Calculating Drug Sensitivity Scores (DSS)

To calculate the DSS we followed our well-detailed method recently published (*52*) which compares the tumor cell survival, measured via bioluminescence imaging, to the health of the OMMCs, measured via PIA. An overall drug sensitivity value resulted from the integrated and meticulous calculations involving eleven combined and weighted parameters: (1) killing at maximum dose (Max Kill), (2) dose required to kill 10% of the tumor (EC10), (3) dose required to kill 25% of the tumor (EC25), (4) dose required to kill 50% of the tumor (EC50), (5) dose required to kill 75% of the tumor (EC75), (6) dose required to kill 90% of the tumor (EC90), (7) slope through the EC50, (8) the area under the curve (AUC), (9) tumor growth acceleration, (10) biphasic killing (rapid killing at low doses and limited additional killing at higher doses), and (11) incomplete kill at the highest dose. Overall DSS from 0 to 100 signify increasing efficacy in tumor kill relative to slice toxicity, while scores from 0 to −100 describe scenarios in which tumors thrive more effectively than OMMCs for a given treatment. Dose-response values were calculated via linear interpolation of raw data, not from best-fit curve equations.

### Patient Ovarian Tumor Preparation for Engraftment onto OMMCs

After consent, 1 to 4g of metastatic ovarian tumor tissues surgically resected from patients at UNC hospitals were placed in sterile Dulbecco’s Modified Eagle Medium/Nutrient Mixture F-12 (DMEM/F-12) from Gibco^TM^ and kept at 4 °C while transferred to the UNC Tissue Procurement Core Facility (TPF). After approval for release, the tumor sample was processed in the Hingtgen laboratory within 1 hour and minced into pieces of about 0.5 mm diameter using a disposable scalpel and PBS. Tumor pieces were stored in a cryogenic vial and frozen in tissue freezing medium (CryoStor CS10) in a FreezeCell™ at −80°C overnight before transfer into liquid nitrogen. At the time of the engraftment onto the OMMCs the thawed tumor cell suspension was passed through a 100 µm cell strainer and the filtered content was collected. Each 50 mg of the tissue collected was transduced by incubating 1ml mCherry-FLuc Lentivirus at 1.5e7 vg/ml with 2µl polybrene for 4h at 37C^°^. After incubation, the tumor tissue was washed with PBS three times and reconstituted in PBS with a final volume of 200 µl. The tumor cell suspension was divided into ∼200 tumor spots in total which makes ∼0.25 mg tissue in 1 µl of PBS/per tumor spot on OMMCs. After tumor spot placement, plates were incubated at 37C with 5% CO2 and 95% humidity. The culture media was changed after 24h and was subsequently changed every 3 days. The initial viability of the patient tissue was measured by BLI on Day 3 for all the bearing-tumor mesenteries before any chemotherapy went into the media and a second BLI was read on Day 6. Data values were normalized to Day 3 for all groups and transformed into survival percentages. Each group was compared to an untreated control group.

### Preparing Human Tumor Tissue for in vitro culture and survival

The frozen tissue was mechanically dissected and filtered into a near single-cell suspension with PBS under sterile conditions as described above then divided into 4 equal portions. Each portion was centrifuged, and the supernatant was removed. They were reconstituted with 4 different growth mediums: (1) DMEM with 20% FBS and 1% PS), (2) MEM with 20% FBS, 1% PS, and insulin human recombinant Zinc (Gibco^TM^)(3) DMEM/F-12 with 20% FBS and 1% PS and (4) McCoy’s 5A (modified) Medium all from Gibco^TM^ with 20% FBS, and 1% PS. All cultures were initiated in a volume of 1 ml per well using 12 well plates (Corning Life Sciences) and incubated at 37°C and 5% CO2. The viable cells and tissue chunks were allowed to settle and attach to the bottom of the plates for 2-3 days. The floating cellular debris was then carefully aspirated, wells were washed with PBS, and 2 ml of fresh medium was added. A potential tumor cell survival was visually checked every two days and also by BLI on Day 6. The culture medium was routinely changed every 3∼5 days.

### Statistical analysis

All statistical tests and sample sizes are included in Figure Legends. All data are shown as mean ± Std. In all cases, the p values are represented as follows: ****p < 0.0001, *** p < 0.001, **p < 0.01, *p <0.05, and not statistically significant when p > 0.05. In all cases, the stated ‘‘n’’ value is either the number of OMMCs, the number of tumor spots placed on the OMMCs, or mice with multiple independent images used to obtain data points for each. Mean values between two groups were compared using t-tests with Welch’s correction when variances were deemed significant by F tests. Mean values between three or more groups were compared to the control by using one-way ANOVA followed by Dunnett’s multiple comparisons test. All statistical analyses were performed using GraphPad Prism (Version 9.1.0). For all quantifications of BLI or FL, the samples being compared were processed in parallel and imaged using the same settings, scale, and laser power.

## Acknowledgments

We would like to thank a local 501c3 organization She Rocks, Research Ovarian Cancer Knowledge Support, for funding this study and to make it possible through the guidance of Dr. Bae-Jump. To all the patients and their families who under consent, generously donated tissue, and the Tissue Procurement Facility at UNC hospitals for obtaining each sample. We would also like to thank the Biomedical Research Imaging Center at UNC; Dr. Satterlee from the Eshleman Institute for Innovation at UNC and Dr. Hingtgen from the Division of Pharmacoengineering and Molecular Pharmaceutics at UNC. Some figures were created using BioRender.com.

## REFERENCES

1. American Cancer Society. Cancer Statistics Center. http://cancerstatisticscenter.cancer.org. Accessed June 03, 2023.

2. National Cancer Institute. SEER stat fact sheets: ovarian cancer. SEER http://seer.cancer.gov/statfacts/html/ovary.html. Accessed June 03, 2023.

3. Shield K, Ackland ML, Ahmed N, Rice GE. Multicellular Spheroids in Ovarian Cancer Metastases: Biology and Pathology. Gynecol Oncol,113(1):143–8 (2009).

4. Raghavan S, Mehta P, Ward MR, Bregenzer ME, Fleck EMA, Tan L, et al. Personalized Medicine-Based Approach to Model Patterns of Chemoresistance and Tumor Recurrence Using Ovarian Cancer Stem Cell Spheroids. Clin Cancer Res, 23(22):6934– 45 (2017).

5. Parashar D, Geethadevi A, Mittal S, et al. Patient-Derived Ovarian Cancer Spheroids Rely on PI3K-AKT Signaling Addiction for Cancer Stemness and Chemoresistance. Cancers (Basel), 14(4):958 (2022).

6. Weroha SJ, Becker MA, Enderica-Gonzalez S, Harrington SC, Oberg AL, Maurer MJ, et al. Tumorgrafts as In Vivo Surrogates for Women With Ovarian Cancer. Clin Cancer Res, 20(5):1288–97 (2014).

7. Kleinmanns K, Bischof K, Anandan S, Popa M, Akslen LA, Fosse V, et al. Cd24-Targeted Fluorescence Imaging in Patient-Derived Xenograft Models of High-Grade Serous Ovarian Carcinoma. EBioMedicine, 56:102782 (2020).

8. Cybula M, Wang L, Wang L, Drumond-Bock AL, Moxley KM, Benbrook DM, Gunderson-Jackson C, Ruiz-Echevarria MJ, Bhattacharya R, Mukherjee P, Bieniasz M. Patient-Derived Xenografts of High-Grade Serous Ovarian Cancer Subtype as a Powerful Tool in Pre-Clinical Research. Cancers (Basel), 13(24):6288 (2021).

9. Nanki Y, Chiyoda T, Hirasawa A, Ookubo A, Itoh M, Ueno M, et al. Patient-derived ovarian Cancer Organoids Capture the Genomic Profiles of Primary Tumours Applicable for Drug Sensitivity and Resistance Testing. Sci Rep, 10(1):12581 (2020).

10. Hill SJ, Decker B, Roberts EA, Horowitz NS, Muto MG, Worley MJ Jr., et al. Prediction of DNA Repair Inhibitor Response in Short-Term Patient-Derived Ovarian Cancer Organoids. Cancer Discov, 8(11):1404–21(2018).

11. Jabs J, Zickgraf FM, Park J, Wagner S, Jiang X, Jechow K, et al. ScreeningDrug Effects in Patient-Derived Cancer Cells Links Organoid Responses to Genome Alterations. Mol Syst Biol, 13(11): 955 (2017).

12. Kopper O, de Witte CJ, Lohmussaar K, Valle-Inclan JE, Hami N, Kester L, et al. An Organoid Platform for Ovarian Cancer Captures Intra- and Interpatient Heterogeneity. Nat Med (2019) 25(5):838–49 (2019).

13. de Witte CJ, Espejo Valle-Inclan J, Hami N, Lõhmussaar K, Kopper O, Vreuls CPH, et al. Patient-Derived Ovarian Cancer Organoids Mimic Clinical Response and Exhibit Heterogeneous Inter- and Intrapatient Drug Responses. Cell Rep, 31(11):107762 (2020).

14. Maru Y, Tanaka N, Itami M, Hippo Y. Efficient Use of Patient-Derived Organoids as a Preclinical Model for Gynecologic Tumors. Gynecol Oncol,154(1):189–98 (2019).

15. Powley IR, Patel M, Miles G, Pringle H, Howells L, Thomas A, et al. Patient-Derived Explants (PDEs) as a Powerful Preclinical Platform for Anti-Cancer Drug and Biomarker Discovery. Br J Cancer,122(6):735–44 (2020).

16. Abreu, S., Silva, F., Mendes, R. et al. Patient-derived ovarian cancer explants: preserved viability and histopathological features in long-term agitation-based cultures. Sci Rep 10, 19462 (2020).

17. Kenny HA, Lal-Nag M, White EA, Shen M, Chiang CY, Mitra AK, et al. Quantitative High Throughput Screening Using a Primary Human Three-Dimensional Organotypic Culture Predicts In Vivo Efficacy. Nat Commun, 6:6220 (2015).

18. Lal-Nag M, McGee L, Guha R, Lengyel E, Kenny HA, Ferrer M. A High-Throughput Screening Model of the Tumor Microenvironment for Ovarian Cancer Cell Growth. SLAS Discov, 22(5):494–506 (2017).

19. Kenny HA, Lal-Nag M, Shen M, Kara B, Nahotko DA, Wroblewski K, et al. Quantitative High-Throughput Screening Using an Organotypic Model Identifies Compounds That Inhibit Ovarian Cancer Metastasis. Mol Cancer Ther, 19(1):52–62 (2020).

20. Watters KM, Bajwa P, Kenny HA. Organotypic 3d Models of the Ovarian Cancer Tumor Microenvironment. Cancers (Basel), 10(8):265 (2018).

21. N. Joshi, D. Liu, K.A. Dickson, D.J. Marsh, C.E. Ford, M.H. Stenzel An organotypic model of high-grade serous ovarian cancer to test the anti-metastatic potential of ROR2 targeted polyion complex nanoparticles J. Mater. Chem. B, 9: 9123–9135 (2021).

22. Davies EJ, Dong M, Gutekunst M, Närhi K, van Zoggel HJ, Blom S, et al. Capturing Complex Tumour Biology In Vitro: Histological and Molecular Characterisation of Precision Cut Slices. Sci Rep 5:17187 (2015).

23. Meijer TG, Naipal KA, Jager A, van Gent DC. Ex Vivo Tumor Culture Systems for Functional Drug Testing and Therapy Response Prediction. Future Sci OA, 3(2):FSO190 (2017).

24. Sivakumar R, Chan M, Shin JS, Nishida-Aoki N, Kenerson HL, Elemento O, et al. Organotypic Tumor Slice Cultures Provide a Versatile Platform for Immuno-Oncology and Drug Discovery. Oncoimmunology, 8(12): e1670019 (2019).

25. Mann B, Zhang X, Bell N, Adefolaju A, Thang M, Dasari R, Kanchi K, Valdivia A, Yang Y, Buckley A, Lettry V, Quinsey C, Rauf Y, Kram D, Cassidy N, Vaziri C, Corcoran DL, Rego S, Jiang Y, Graves LM, Dunn D, Floyd S, Baldwin A, Hingtgen S, Satterlee AB. A living ex vivo platform for functional, personalized brain cancer diagnosis. Cell Rep Med. 4(6):101042 (2023).

26. Nougaret S, Addley HC, Colombo PE, Fujii S, Al Sharif SS, Tirumani SH, Jardon K, Sala E, Reinhold C. Ovarian carcinomatosis: how the radiologist can help plan the surgical approach. Radiographics. 32(6):1775–803 (2012).

27. Anderson CR, Ponce AM, Price RJ. Immunohistochemical identification of an extracellular matrix scaffold that microguides capillary sprouting in vivo. J Histochem Cytochem. 52(8):1063–72 (2004).

28. Zhang N, Kim SH, Gainullina A, Erlich EC, Onufer EJ, Kim J, Czepielewski RS, Helmink BA, Dominguez JR, Saunders BT, Ding J, Williams JW, Jiang JX, Segal BH, Zinselmeyer BH, Randolph GJ, Kim KW. LYVE1+ macrophages of murine peritoneal mesothelium promote omentum-independent ovarian tumor growth. J Exp Med. 218(12):e20210924 (2021)

29. Sandhu SK, Schelman WR, Wilding G, Moreno V, Baird RD, Miranda S, Hylands L, Riisnaes R, Forster M, Omlin A, Kreischer N, Thway K, Gevensleben H, Sun L, Loughney J, Chatterjee M, Toniatti C, Carpenter CL, Iannone R, Kaye SB, de Bono JS, Wenham RM. The poly(ADP-ribose) polymerase inhibitor niraparib (MK4827) in BRCA mutation carriers and patients with sporadic cancer: a phase 1 dose-escalation trial. Lancet Oncol.14(9):882–92 (2013).

30. Gelmon KA, Tischkowitz M, Mackay H, Swenerton K, Robidoux A, Tonkin K, Hirte H, Huntsman D, Clemons M, Gilks B, Yerushalmi R, Macpherson E, Carmichael J, Oza A. Olaparib in patients with recurrent high-grade serous or poorly differentiated ovarian carcinoma or triple-negative breast cancer: a phase 2, multicentre, open-label, non-randomised study. Lancet Oncol,12(9):852–61(2011).

31. Tutt A, Robson M, Garber JE, Domchek SM, Audeh MW, Weitzel JN, Friedlander M, Arun B, Loman N, Schmutzler RK, Wardley A, Mitchell G, Earl H, Wickens M, Carmichael J. Oral poly(ADP-ribose) polymerase inhibitor olaparib in patients with BRCA1 or BRCA2 mutations and advanced breast cancer: a proof-of-concept trial. Lancet, 376(9737):235–44 (2010).

32. Mirza MR, Pignata S, Ledermann JA. Latest clinical evidence and further development of PARP inhibitors in ovarian cancer. Ann Oncol, 29(6):1366–1376, (2018).

33. Kristeleit R, Lisyanskaya A, Fedenko A, Dvorkin M, de Melo AC, Shparyk Y, Rakhmatullina I, Bondarenko I, Colombo N, Svintsitskiy V, Biela L, Nechaeva M, Lorusso D, Scambia G, Cibula D, Póka R, Oaknin A, Safra T, Mackowiak-Matejczyk B, Ma L, Thomas D, Lin KK, McLachlan K, Goble S, Oza AM. Rucaparib versus standard-of-care chemotherapy in patients with relapsed ovarian cancer and a deleterious BRCA1 or BRCA2 mutation (ARIEL4): an international, open-label, randomised, phase 3 trial. Lancet Oncol, 23(4):465–478 (2022).

34. Penson RT, Valencia RV, Cibula D, Colombo N, Leath CA 3rd, Bidziński M, Kim JW, Nam JH, Madry R, Hernández C, Mora PAR, Ryu SY, Milenkova T, Lowe ES, Barker L, Scambia G. Olaparib Versus Nonplatinum Chemotherapy in Patients With Platinum-Sensitive Relapsed Ovarian Cancer and a Germline BRCA1/2 Mutation (SOLO3): A Randomized Phase III Trial. J Clin Oncol, 38(11):1164–1174 (2020).

35. Kim, YN., Park, B., Kim, J.W. et al. Triplet maintenance therapy of olaparib, pembrolizumab and bevacizumab in women with *BRCA* wild-type, platinum-sensitive recurrent ovarian cancer: the multicenter, single-arm phase II study OPEB-01/APGOT-OV4. Nat Commun 14, 5476 (2023).

36. Genomics of Drug Sensitivity in Cancer. https://www.cancerrxgene.org/about. Accessed, October, 2023.

37. Eric J Devor, Jace R Lapierre, David P Bender. ES-2 Ovarian Cancer Cells Present a Genomic Profile Inconsistent with their Reported History. Obstet Gynecol Res, 4 (4): 233–238 (2021).

38. Smeby J, Kryeziu K, Berg KCG, Eilertsen IA, Eide PW, Johannessen B, Guren MG, Nesbakken A, Bruun J, Lothe RA, Sveen A. Molecular correlates of sensitivity to PARP inhibition beyond homologous recombination deficiency in pre-clinical models of colorectal cancer point to wild-type TP53 activity. EBioMedicine, 59:102923 (2020).

39. Shen, J.; Peng, Y.; Wei, L.; Zhang, W.; Yang, L.; Lan, L.; Kapoor, P.; Ju, Z.; Mo, Q.; Shih Ie, M.;, et al. ARID1A Deficiency Impairs the DNA Damage Checkpoint and Sensitizes Cells to PARP Inhibitors. Cancer Discov. 5, 752–767 (2015).

40. Kuroda T, Kohno T. Precision Medicine for Ovarian Clear Cell Carcinoma Based on Gene Alterations. Int J Clin Oncol. 25:419–24 (2020).

41. Saito R, Kuroda T, Yoshida H, Sudo K, Saito M, Tanabe H, Takano H, Yamada K, Kiyokawa T, Yonemori K, Kato T, Okamoto A, Kohno T. Genetic characteristics of platinum-sensitive ovarian clear cell carcinoma. Jpn J Clin Oncol. 53(9):781–790, (2023).

42. Jang JYA, Yanaihara N, Pujade-Lauraine E, Mikami Y, Oda K, Bookman M, Ledermann J, Shimada M, Kiyokawa T, Kim BG, Matsumura N, Kaku T, Kuroda T, Nagayoshi Y, Kawabata A, Iida Y, Kim JW, Quinn M, Okamoto A. Update on rare epithelial ovarian cancers: based on the Rare Ovarian Tumors Young Investigator Conference. J Gynecol Oncol. 28(4): e54 (2017).

43. Takahashi K, Takenaka M, Okamoto A, Bowtell DDL, Kohno T. Treatment Strategies for ARID1A-Deficient Ovarian Clear Cell Carcinoma. Cancers (Basel).13(8):1769 (2017).

