## Supplementary figures and images for "A Novel ex-vivo platform for personalized treatment in metastatic ovarian cancer"

### Supplemental Figures

#### Slide 1
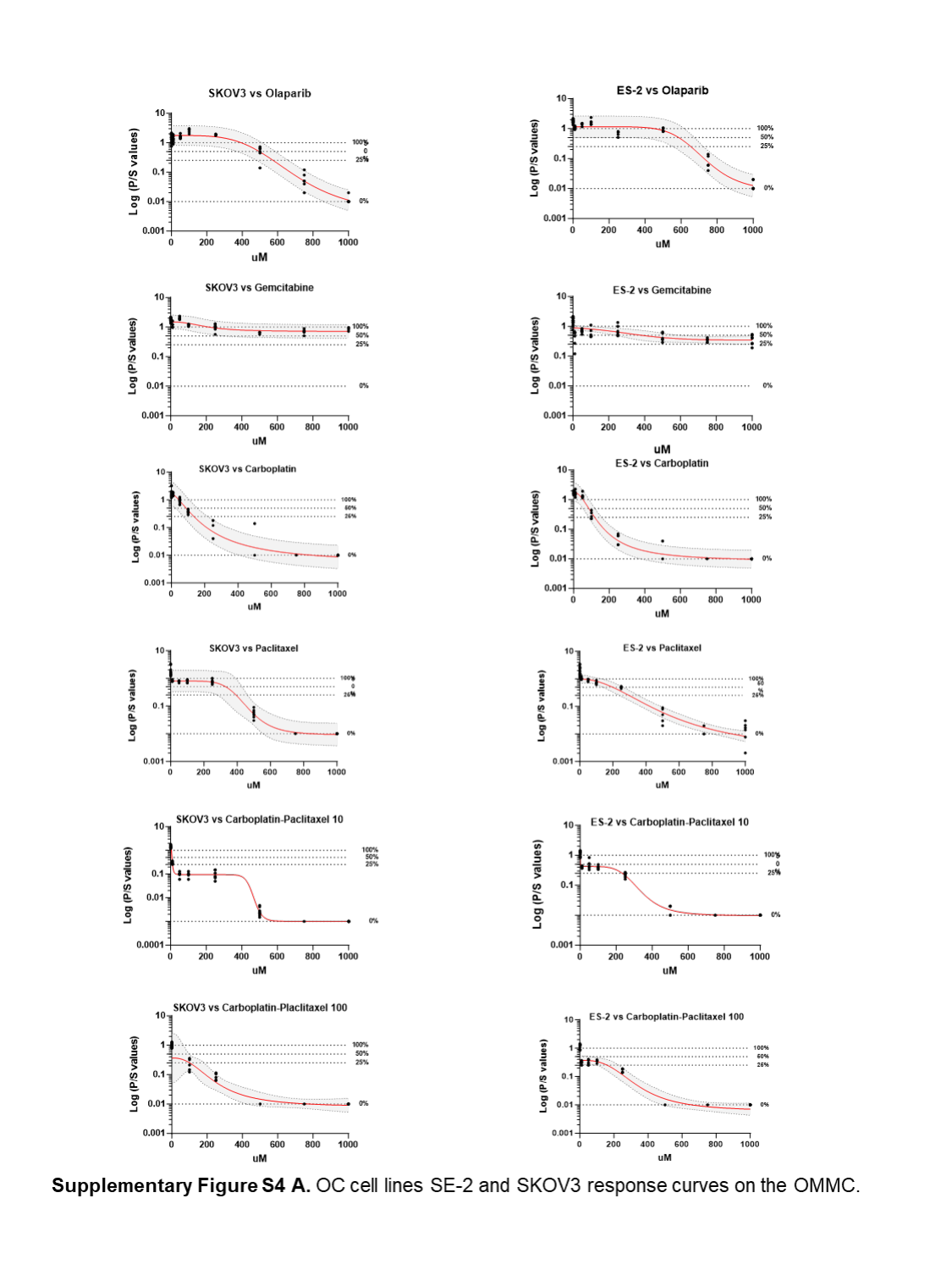

#### Slide 2
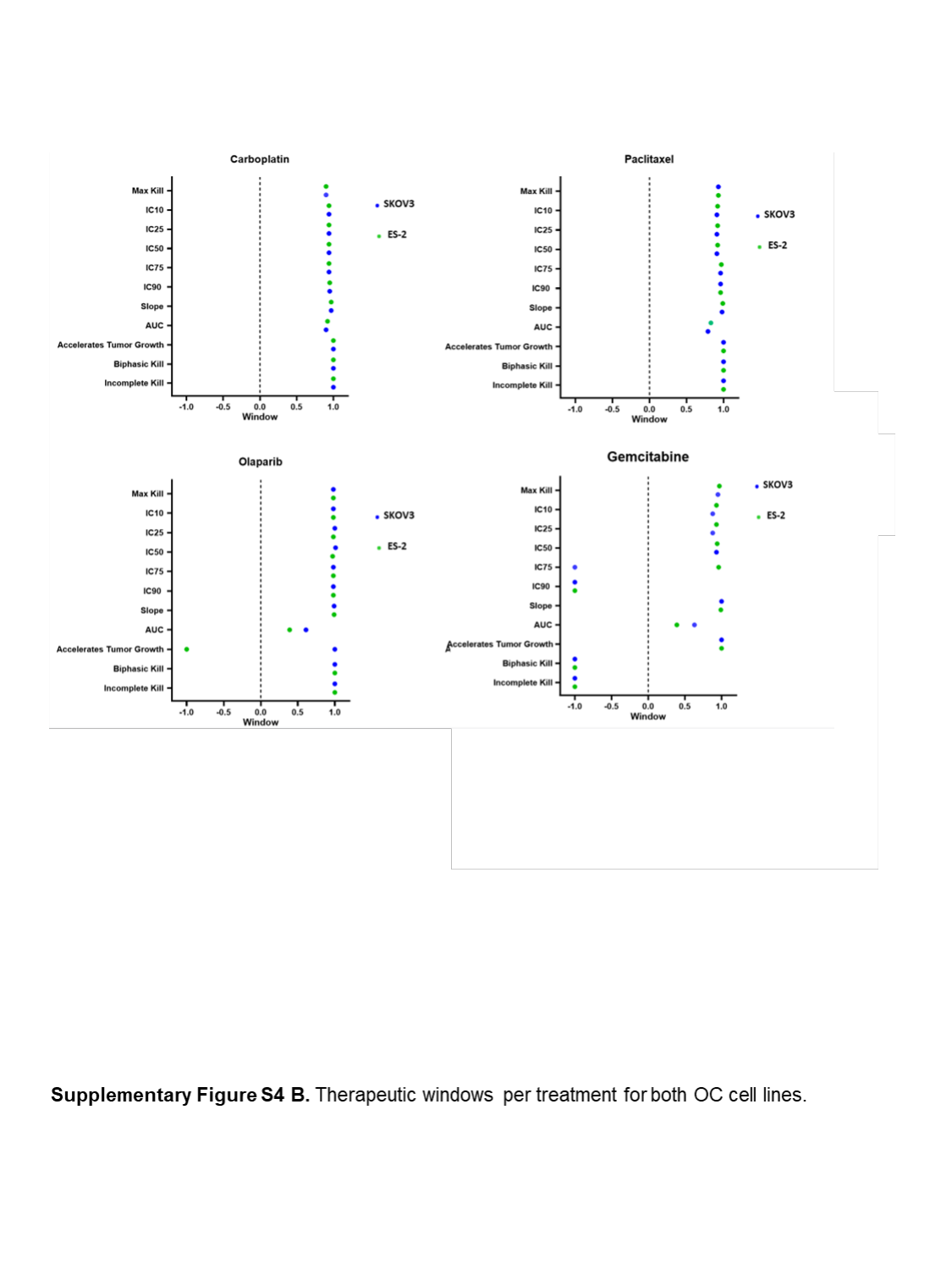

#### Slide 3
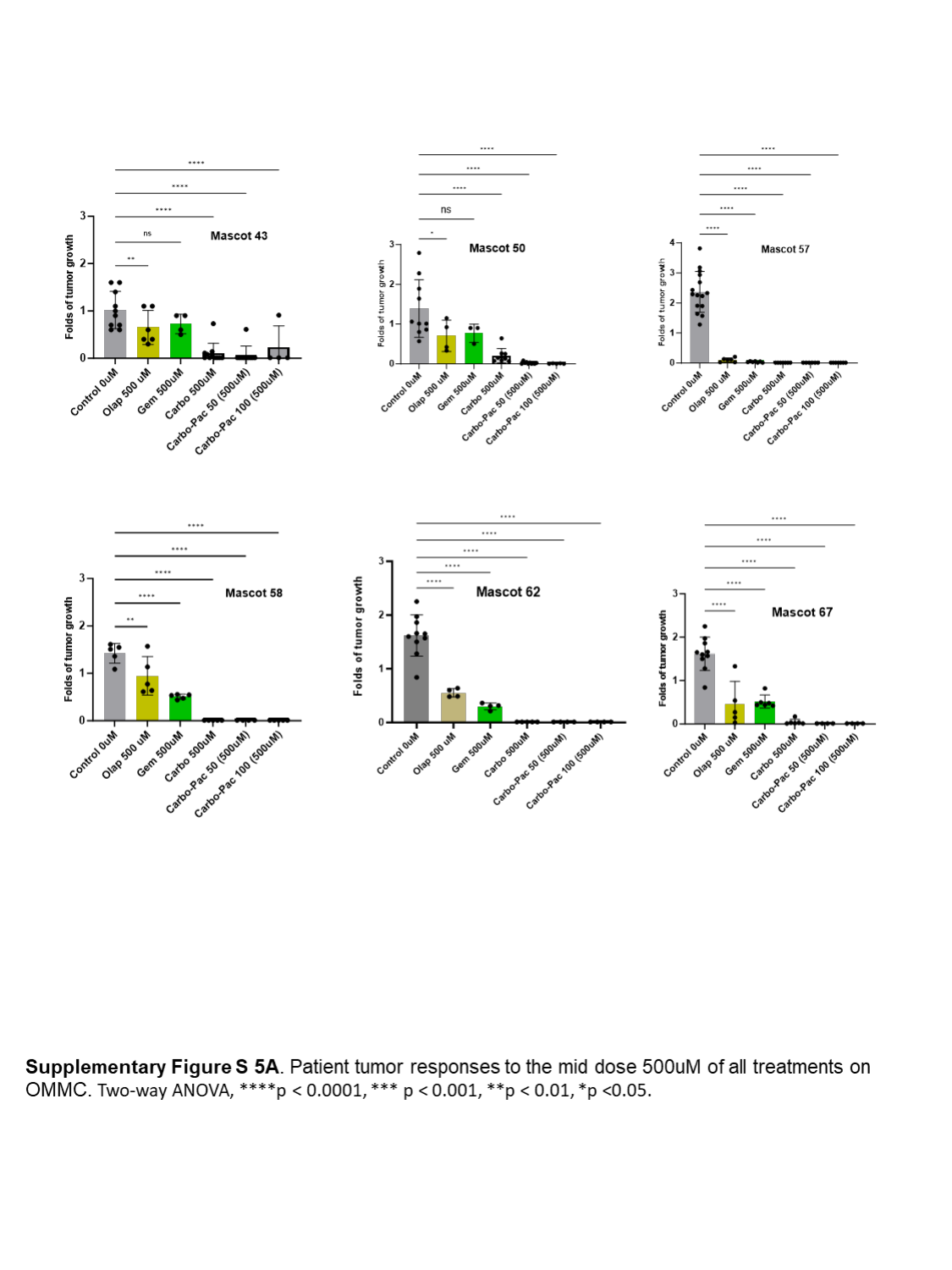

#### Slide 4
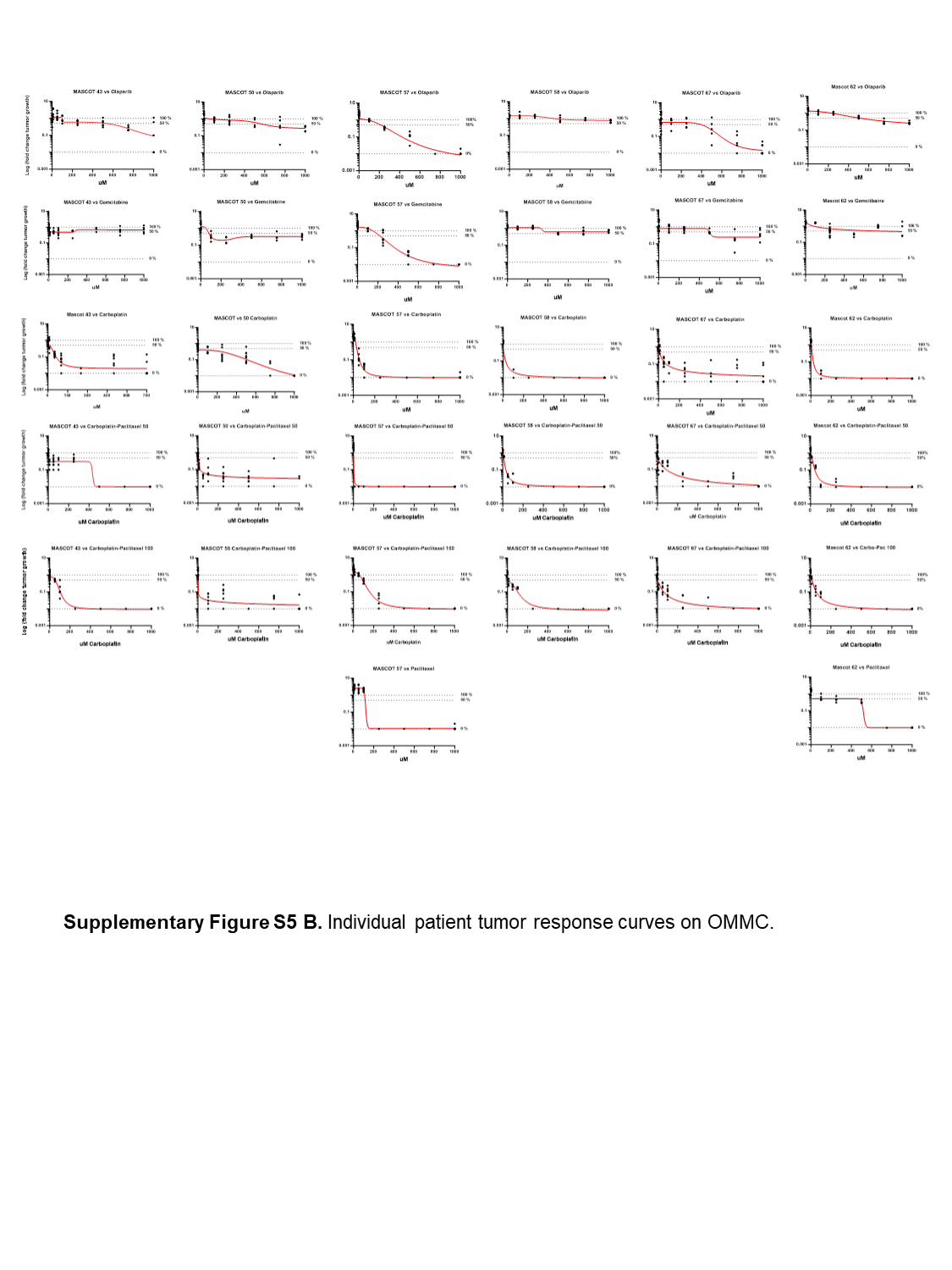
